# Chlorpyrifos and tamoxifen co-pretreatment promotes stem-like phenotype and upregulation of anti-estrogen therapy resistance markers in ERα breast cancer cells

**DOI:** 10.1101/2025.08.30.673130

**Authors:** Marianela Lasagna, Lucia Enriquez, Mariana Mardirosian, Rocio Natalia Cardozo, Gabriela Martín, Pablo Power, Mariel Nuñez, Claudia Cocca

**Affiliations:** Instituto de Química y Fisicoquímica Biológicas "Prof. Alejandro C. Paladini" (IQUIFIB) UBA-CONICET, Buenos Aires, Argentina; Laboratorio de Radioisótopos, Cátedra de Física, Facultad de Farmacia y Bioquímica, Universidad de Buenos Aires, Buenos Aires, Argentina; Departamento de Microbiología, Inmunología y Biotecnología, Facultad de Farmacia y Bioquímica, Universidad de Buenos Aires, Buenos Aires, Argentina

**Author notes:** Corresponding author: Marianela Lasagna. Electronic address. Marianela Lasagna, Lucia Enriquez, Mariana Mardirosian Roció Natalia Cardozo Gabriela Martin Pablo Power, Mariel Núñez Claudia Cocca.

**Keywords:** Breast Cancer, Endocrine Therapy Resistance, Chlorpyrifos, Histone Deacetylase 1, Stemness Phenotype

## Abstract

Endocrine therapy (ET) is the standard first-line treatment for hormone receptor positive breast cancer but about 30% of patients develop resistance. Chlorpyrifos (CPF), is an endocrine disruptor pesticide that promotes a stemness phenotype and triggers molecular mechanisms involved in ET resistance. The goal of this study is to assess whether CPF induces tamoxifen (TAM) resistance by modulating estrogen receptor alpha (REα) expression through histone deacetylase 1 (HDAC1) regulation. We analyzed public databases of breast cancer patients and performed *in vitro* experiments using the MCF-7 cell line. We evaluated whether the co-pretreatment with CPF and TAM (CPF+TAM) modulates stemness and ET resistance markers. We found that CPF+TAM selects cells that exhibit increased clonogenicity, enhanced mammosphere-forming ability, and upregulates the stemness markers CD44, NANOG and POU5F1 while decreasing CD24. This selected cell subpopulation shows higher levels of mRNA and elevated protein expression of REα and HDAC1. Finally, we identified a resistance and stemness marker signature induced by CPF+TAM that closely resembles the profile observed in the dataset of patients who acquired tamoxifen resistance. Our findings show that CPF promotes an undifferentiated basal-like cell phenotype that contributes to TAM resistance, reinforcing the need for global restrictions to safeguard public health.

## 1. Introduction

Despite the implementation of numerous prevention strategies and advances in treatment, breast cancer remains the most common cancer in women worldwide (Nolan et al., 2023). More than two million new cases of breast cancer (BC) were diagnosed worldwide in 2022 (Sung et al., 2021). Among them, 70-80% of patients present with hormone receptor (HR) positive breast cancer (Estrogen Receptor α (ERα) and Progesterone Receptor (PR)), which according to the intrinsic molecular subtype classification of PAM50 are defined as luminal subtypes A and B (Al-thoubaity, 2020). In these subtypes, oestrogens (E2) play a key role in tumour initiation and progression (Gu et al., 2010). Regardless of the fact that E2 has said BC-promoting effects, HR-positive breast carcinomas, when compared to their ER-negative counterparts, display slower proliferation, lower levels of division markers such as Ki67, less frequency of metastasis, and are often responsive to endocrine therapy (ET) (Li et al., 2021). This apparent paradox denotes that the ER network has a positive value even in a transformed environment (Sukocheva et al., 2022).

Proliferation and survival of tumor cells depends on ERα activation, which upon binding to endogenous estrogens dimerises and translocates to the nucleus. ER dimers bind to coactivators to form a transcriptionally active complex promoting the transcription of pro-survival genes (genomic regulation). Additionally, the binding of ER to endogenous ligands can lead to the activation of different signaling pathways through non-genomic regulation (Hanker et al., 2020).

Due to the strong dependence of breast tumorigenesis on the estrogen-RE axis, ET, such as selective ER modulators (SERMs), selective ER downregulators (SERDs), and aromatase inhibitors (AIs) remain the first-line treatment in patients with HR-positive breast cancer (Aggelis & Johnston, 2019). Unfortunately, resistance to ET is seen in 30% of early breast cancer cases and develops in almost all patients with advanced disease (Pan et al., 2017).

Loss of ER expression is known to occur in approximately 20% of endocrine-resistant breast cancers (Schrijver et al., 2018). Endocrine resistance is usually due to ligand-independent ER reactivation. This can occur due to mutations in the ER leading to increased ER activity (Z. Li et al., 2022) altered interactions of the ER with coactivators/corepressors or through compensatory interaction between the ER and growth factor receptors and oncogenic signaling pathways (Ma et al., 2015).

The aforementioned ER status is under efficient regulation by specific epigenetic mechanisms, which include hypermethylation of CpG islands within ER promoters, increased histone deacetylase (HDACs) activity in the ER promoter, and/or translational repression by miRNAs. Additionally, high methylation of the ERα gene (ESR1) promoter by DNA methyltransferases was linked to poor prognosis, and its profile was suggested as a tool to predict response to hormonal and non-hormonal therapy in BCs(Sukocheva et al., 2022).

The aforementioned HDACs form multi-unit protein complexes and function as transcriptional repressors by removing acetyl groups from histone lysine residues (Seto & Yoshida, 2014). In this sense, HDAC1 is known to interact with ERα and suppress the transcriptional activity of ERα by increasing the level of chromatin condensation (Dimitrakopoulos et al., 2021). Additionally, Tamoxifen (TAM) binding to ER promotes interaction with corepressors such as LCOR and NCOR1/2, recruits epigenetic repressors such as HDACs and the NuRD complex to mediate the removal of active epigenetic marks and represses ER transcriptional activity. These regulation mechanisms are closely related to the development of resistance to ET (Garcia-Martinez et al., 2021).

In addition to the previously mentioned mechanisms, recent studies propose that acquired resistance to ET may be due to a more dynamic process involving phenotypic plasticity and tumor heterogeneity(Menon et al., 2020). Cancer stem cells (CSCs) are characterized by two fundamental properties, namely their ability to self-propagate and to restore the original tumor with phenotypic heterogeneity (Valent et al., 2012). Another characteristic of CSCs is the development of resistance to unfavorable microenvironments, such as those generated by ET. One plausible explanation is that these cells adopt a slow-cycling or semi-quiescent phenotype, which renders them more resilient to various stressors. Subsequently, they can swiftly transition to a proliferative state when environmental conditions become more conducive, thereby enhancing their tumorigenic potential (Recasens & Munoz, 2019).

It has long been known that mammary CSCs express receptors for different types of hormones and can be affected by the action of estrogen and progesterone (Zheng & Karnoub, 2021). However, the effects of estrogen on the mammary CSC subpopulation are not fully understood and remain under review (Simões et al., 2011). While some authors propose that estrogens promote the expansion of CSCs through both genomic and non-genomic pathways (Zhou et al., 2015), others argue that it is the inhibition of estrogen signaling that is indeed associated with CSC expansion since estrogens were shown to reduce CSC self-renewal in the MCF-7 cell line by down-regulating the expression of pluripotency genes such as NANOG, OCT4 and SOX2 (Simões & Vivanco, 2011).

ET used in breast cancer often target differentiated proliferative cells, but have a controversial effect on CSCs, with an increase in this subpopulation even observed after anti-estrogen treatment(Garcia- Recio et al., 2023). In addition, treatment with TAM increased both the number of mammospheres and the expression of pluripotency markers such as NANOG, OCT4 and SOX2 in cancer cells (Piva et al., 2014; Simões et al., 2011). It was demonstrated that MCF-7 mammospheres are resistant to high doses of TAM(Cariati et al., 2008).

Endocrine-disrupting chemicals (EDCs) that mimic estradiol, and affect its pathways either by binding to ER, modifying estradiol metabolism, regarding its biosynthesis or degradation, or by promoting its genomic and non-genomic activation. Given the well known carcinogenic effect that estrogens have on breast epithelial cells, there has been great efforts in studying the impact of EDs on the different steps of breast cancer, including their involvement in the development of resistance to ET(Koual et al., 2020; Rodgers et al., 2018). The organophosphate chlorpyrifos (CPF) is one of the most widely used pesticides in Argentina and the world, to which we are ubiquitously exposed as it is present in fruits and vegetables. It is considered an EDCs, being able to mimic hormonal actions both in vitro and in vivo models. Furthermore, it promotes migration and invasion as well as epithelial- mesenchymal transition (EMT) in breast cancer cells in vitro(Lasagna et al., 2020, 2022). In addition, we recently demonstrated that CPF promotes lung metastasis in exposed mice (Lasagna et al., 2025) and modulates the expression of MET and cancer stem cell (CSC) markers in our tumor model.

In this context, as an initial approach, publicly available datasets of breast cancer patients were analysed to identify whether stemness markers and resistance to therapy correlate with patient prognosis and disease progression. Subsequently, it was determined whether exposure to CPF and TAM promotes tumor heterogeneity by inducing cell proliferation with a stemness phenotype, which could be mediated by the participation of HDAC1 and RE, leading to tumor progression toward more aggressive behavior and resistance to antiestrogen therapy.

## 2. Materials and methods

### 2.1 Analysis of Publicly Available Datasets on TAM Resistance in Breast Cancer

#### 2.1.1 Gene expression data collection

Gene expression profiles were downloaded from the NCBI Gene Expression Omnibus (GEO) database (https://www.ncbi.nlm.nih.gov/gds). Initially we performed the analysis of the GSE241654 dataset comparing the expression profiles of MCF-7 wt and TAM-resistant MCF-7 cells. Subsequently, we analysed datasets containing gene expression profiles from clinical samples of TAM-treated (GSE6532, GSE9195, GSE17705, GSE12093) and TAM-untreated (GSE2034, GSE7390 y GSE6532) RE+ breast cancer patients (Simões et al., 2015). To make all microarrays comparable we use Robust Multiarray Average (RMA) which method of the Affy packages (v1.82.0) performs background fluorescence correction, normalisation, summation and estimation of expression levels in log2. Probe annotation was performed using the annotation package provided by the manufacturer: “hgu133A.db” (Carlson, 2023), for the GSE6532, GSE9195 series and "hgu133plus2. db" (Carlson, 2025) for the GSE241654, GSE6532, GSE17705, GSE 12093, GSE2034, GSE7390 and GSE6532 series.

#### 2.1.2 GO annotation and KEGG enrichment analysis in MCF-7 wt and TAM-resistant MCF-7 cells

Differentially expressed genes (DEGs) were identified in Dataset GSE241654 using the exploratory fold change (FC) method. For the selection of DEGs, we used the cut-off criterion log2 Fold Change>1 for the FC method. Then, we performed the Gene Ontology (GO) annotation and Kyoto Encyclopedia of Genes and Genomes (KEGG) pathway enrichment analysis. The GO analysis includes three categories: biological process (BP), cellular component (CC) and molecular function (MF). Finally, a bubble chart was plotted using the Bioinformatics (www.bioinformatics.com.cn) free online platform for bioinformatics-related data analysis.

#### 2.1.3 Expression profile analysis, clinical outcomes and markers correlation

We compared the expression levels of genes of interest related to the development of TAM resistance: ESR1, HDAC1 and NCOR2 and to the stemness phenotype: CD44, CD24, NANOG, SOX2 and POU5F1 in TAM-treated and TAM-untreated patients. Patients in the cohort were divided according to HDAC1 expression levels: low HDAC1 (<25th percentile), medium HDAC1 (25th-75th percentile), high HDAC1 (>75th percentile). Information on clinicopathological characteristics was retrieved and the diseases-free survival of patients with high and low HDAC1 in TAM-treated and TAM-untreated groups was compared and visualised with the Kaplan-Meier method. Additionally, the correlation of the aforementioned genes of interest in each of the TAM-treated and TAM-untreated groups was assessed. Finally, we determined whether and when patients had relapsed, configuring 3 groups of analysis: patients without relapse, primary endocrine resistance (relapse before 2 years of treatment) or secondary endocrine resistance (relapse after 2 years of treatment) (Cardoso et al., 2024). The mean expression levels of HDAC1 and ESR1 were compared between the groups, as well as the correlation of the genes of interest.

### 2.2 Cell culture and exposure models

The estrogen-dependent MCF-7 breast cancer cells were cultured and we performed mammosphere formation and clonogenic assays as described previously (Lasagna et al., 2022; Ventura et al., 2015). We carried out three different exposure models.

#### CPF pretreated model

This model consists of pre-treatment with vehicle or CPF (0.05 or 50) for 72 h. Cells are then trypsinised and seeded on adherent (clonogenic assay) or ultra-low adherent (mammosphere formation assay) plates. Then added TAM (0.1 μM) to the culture medium.

#### CPF+TAM concurrently co-pretreated model

This model consisted in exposing cells concurrently to either vehicle or CPF (0.05 μM and 50 μM) and TAM (0.2 μM and 0.5 μM) for 72 h. Then, cells were trypsinised and seeded on adherent or ultra-low adherent plates.

### 2.3 TAM concentration-response curve

We constructed concentration-response calibration curves to assess the effects of TAM citrate (gift from Gador Laboratories SA, Buenos Aires, Argentina) on clonogenicity and mammosphere formation assay. Initially, TAM was added to the culture medium and used in the “CPF pretreated model”. Furthermore, since the “CPF+TAM concurrently co-pretreated model” was based on pre-treatment of MCF-7 cells for 72 h with CPF, we decided to perform calibration curves with MCF-7 cells pre-treated for 72 h at different concentrations of TAM to evaluate clonogenicity and mammosphere formation.

### 2.4 Molecular Docking

The structural model of the human estrogen receptor-alpha (hERα) ligand binding domain (LBD) was obtained from the PDB database (https://www.rcsb.org/structure/1A52), and used as a template for the performance of molecular docking assays. The spatial coordinates of the ligands estradiol (E2), CPF, and TAM were retrieved from the ZINC (https://zinc15.docking.org/) database, and the structures were minimized in Avogadro v1. The molecular docking between the receptor LBD and the 3 ligands was carried out by Autodock Vina, using the virtual environment of the Yasara software. PyMOL v3.0.0 was used to visualize the molecular interactions and for preparing the figures.

### 2.5 Quantitative real-time PCR analysis (RT-qPCR)

Quick-Zol (Kalium Technologies, Argentina) was used for 5 minutes to isolate RNA from MCF-7 cells cultured in monolayer or as mammospheres to allow complete dissociation of proteins, as described previously (Lasagna, 2025). Total RNA was quantified and RNA quality was determined by measuring optical densities at 260 and 280 nm using the BioTek Take3 device (Agilent, USA). cDNA was synthesized from the RNA template using MMLV reverse transcriptase (Promega Corporation, USA) according to the manufacturer’s instructions. Relative expression levels of ESR1, HDAC1, NCOR2, CD44, CD24, NANOG, SOX2, POU5F1 and β-actin were analyzed following the standard qRT-PCR protocol with the Step One System (Applied Biosystems, USA). The primer pairs used are shown in ***Table 1***. The ΔΔCt method was used for relative mRNA quantification using β-actin as a reference gene.

**Table 1.**
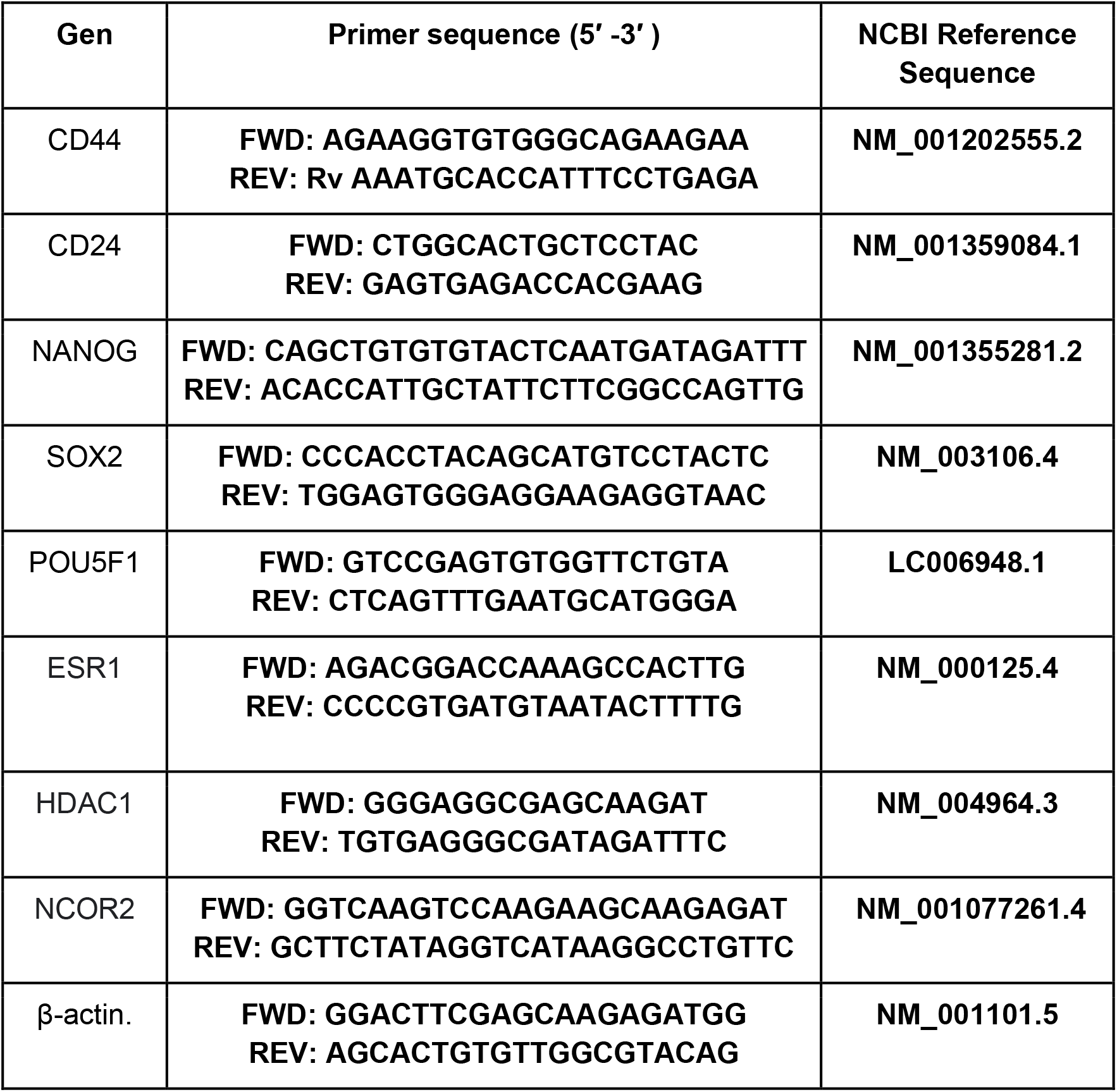

### 2.6 Immunofluorescence

MCF-7 cells were seeded onto coverslips and pretreated with CPF (0.05 and 50 μM) or vehicle in the presence or absence of 0.5 μM TAM for 72 h. Then, they were fixed with 80% methanol and exposed to mouse monoclonal anti-HDAC1 primary antibody (1:200, Sigma Aldrich, USA, donated by Dr. Candela R. Gonzalez, ININFA-UBA-CONICET) and rabbit monoclonal anti-RE primary antibody (1:100, Santa Cruz Biotechnology, Inc, USA) overnight in a humid chamber. Then 2 washes were performed with PBS and incubated for 1 hour with Alexa 567 anti-mouse antibody 1:400 (Red) and Alexa 488-conjugated anti-rabbit antibody 1:400 (Green). Hoechst was used for nuclei staining. The coverslips were mounted with Mowiol. The images were obtained with Zeiss LSM880 confocal microscope, with a Plan-Apochromat 63x objective, Numerical Aperture 1.4. The lasers used were: Multi Argon 458 / 488 / 514 25mW, Diode laser 405 nm, 30 mW, laser class 3B and He Neon GREEN 561 nm, 1.5 mW, laser class 3B. The images were acquired with ZEN BLACK software. Images were taken from 5 random fields. Each of them was Z Stacked at 0.6 μm. For the quantification of fluorescence intensity, a Zproject of each of the stacks was performed. The mean fluorescence intensity (MFI) was determined for each of the nuclei, cytoplasm and/or membrane present in the image using Fiji ImageJ software.

### 2.7 Flow cytometry analysis

For flow cytometric analysis, MCF-7 cells treated with vehicle or CPF (0.05-50 µM) in the presence or absence of 0.05 TAM were trypsinised (trypsin 0.25 %) and fixed with 80% methanol. They were then washed with phosphate buffered saline (PBS) to remove excess methanol. Then a blocking solution (PBS + 10 % SBF) was added for 1 h. Subsequently, anti-mouse CD44 (1:100, Santa Cruz, USA) was added for 1 h, washed with PBS, and then secondary anti-mouse IgG Alexa Fluor 488 (1:400, Invitrogen, USA) and anti-CD24-PE-Cy7 (1:100, Invitrogen, USA, donated by Dr. Karina A Gomez, INGEBI-CONICET) were added for 1 h at room temperature. Finally, samples were washed with PBS and resuspended in 200 μl of PBS. Flow cytometric analysis was performed on the FACSCanto cytometer (BD Bioscience). The percentage of CD24+/CD44+ subpopulations was calculated by the flow cytometry analysis. For quantification of CD24 and CD44 expression levels, we used the MFI obtained with FlowJo software.

### 2.8 Statistical analysis

Data analysis was conducted using GraphPad Prism version 8.0.1 (GraphPad Software Inc., USA). The Shapiro–Wilk test was employed to evaluate the normality of data distributions. Specific statistical tests are detailed in the respective figure legends. A *p*-value<0.05 was considered indicative of statistical significance.

## 3. Results

### 3.1 HDAC1, ESR1 and stemness-related genes are key modulators of the TAM response in ER+ breast cancer

A dot plot summarizing the enriched Gene Ontology (GO) terms is shown (**Fig. 1A**). In relation to BP we observed a significant enrichment in response to TGF-β and transmembrane receptor serine/threonine protein kinase signalling. Additionally, we observed regulation of epithelial cell proliferation as well as epithelial to mesenchymal transition, processes closely related to the response to steroid hormone. Examination of CC reveals a marked enrichment in alterations of cell motility, cell morphology as well as interaction with collagen-rich extracellular. In MF we observed enrichment in growth factor binding and DNA-binding transcription factors. Complementarily, the rest of the terms are closely related to those observed in the CC associated with cell motility. Finally, seven KEGG pathways most significantly enriched in TAM-resistant MCF-7 cells, namely Wnt, PI3K-AKT, MAPK, stem cell pluripotency-associated, estrogen signalling, Proteoglycan and TGF-β (**Fig. 1B**).

**Fig 1.**
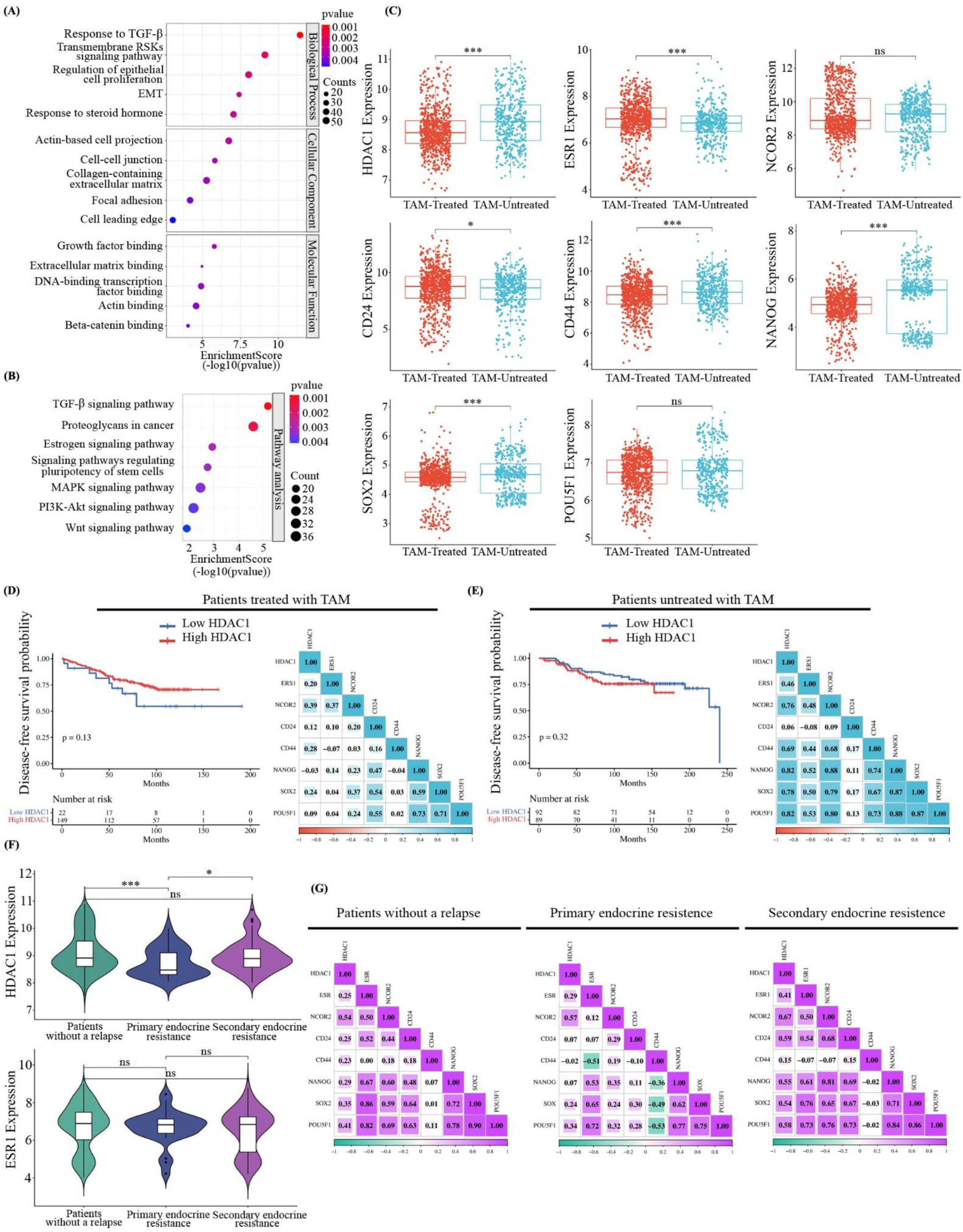
HDAC1 expression and stemness markers correlations in TAM-Treated breast cancer patients. **(A-B)-** Dot plot of Enriched GO and Pathway Enrichment Analysis. Differentially expressed genes in TAM- resistant MCF-7 vs. MCF-7 Wt cells are shown. The color of each dot encodes the statistical significance of the enrichment, expressed as the −log 10 (*p-value*), with warmer colors (towards red) indicating higher significance (lower p-values). The size of each dot is proportional to the number of genes from the dataset that are annotated to that particular GO term, as indicated by the "count" legend. **(C)-** Comparison of HDAC1, ESR1, NCOR2, CD24, CD44, NANOG, SOX2, POU5F1 expression levels between TAM-treated (n=778) and TAM-untreated (n=429) ER+ breast cancer patients. Wilcox test was performed (\**p*<0,05, \*\**p*<0,01, \*\*\**p*<0,001). **(D-E)-** Diseases-free survival in TAM-treated (n=171) and untreated (n=292) patients with low HDAC1 and high HDAC1 expression levels. Significance tested using two-sided t-test. Correlation matrix plot with lower triangle color intensity and size of the square proportional to the correlation coefficients. The color scale bar ranged from −1.0 (red) to 1.0 (light blue). The size and color of the circles indicate the magnitude of the correlation between parameters. Red and light blue, respectively, denote negative and positive correlations. **(F)-** Violin plots illustrating the distribution of HDAC and ESR1 expression levels across three group of ER+ breast cancer patients: those without relapse (n=242), patients with primary endocrine resistance (n=26), and patients with secondary endocrine resistance (n=77). (\**p*<0.05, \*\*\**p*<0.001. One-way ANOVA and Tukey’s Comparison post hoc test). **(G)-** Correlation matrix plot with lower triangle color intensity and size of the square proportional to the correlation coefficients. The color scale bar ranged from −1.0 (light green) to 1.0 (purple). The size and color of the circles indicate the magnitude of the correlation between parameters. Purple and light green, respectively, denote negative and positive correlations.

We also compared expression levels of HDAC1, ESR1 and NCOR2 in patients with ER+ BC treated with TAM against those who were not administered the drug (**Fig. 1C**). We found that the patients receiving ET showed lower levels of HDAC1 (*p*<0.001), as well as higher levels of ESR1 (*p*<0.001) when contrasted against those not treated with TAM. When analyzing the expression of different markers of CSC and indifferentiation, we observed that TAM-treated patients displayed lower levels of CD44 (*p*<0.01), NANOG (*p*<0.001) and Sox2 (*p*<0.001) and higher levels of CD24 (*p*<0.05), while not showing alterations in POU5F1 levels.

Subsequently we evaluated the disease-free survival probability according to the levels of HDAC1 and correlation between the genes of interest mentioned above in TAM-treated and TAM-untreated groups. We found that in the groups of TAM treated patients, those with low HDAC1 levels showed a tendency to relapse earlier than those with high HDAC1 levels (*p*<0.13). Additionally, we found a positive correlation between NCOR2 with HDAC1, ESR and SOX2 (**Fig. 1D**).

In the group that was not treated with TAM, no difference in the disease-free survival probability was observed between the low HDAC1 and high HDAC1 group (*p*<0.35). In this group we observed that HDAC1 positively correlates with ESR1, NCOR2, CD44, NANOG, SOX2, POU5F1. ERS1 positively correlates with NCOR2, CD44, NANOG, SOX2, POU5F1, and NCOR2 positively correlates with CD44, NANOG, SOX2, POU5F1 (**Fig. 1E**).

Furthermore, we evaluated the expression levels of HDAC1 and ESR1 in patients who underwent ET, classifying them according to whether or not they had relapsed (**Fig. 1G**). Patients with primary resistance to TAM showed lower levels of HDAC1 than those who had not relapsed (*p*<0.01) as well as to those with secondary resistance to TAM (*p*<0.05). In contrast, no variations in ESR1 levels were observed among the different groups.

The patients treated with TAM who did not relapse, HDAC1 showed a positive correlation with NCOR2, and POU5F1; ESR1 with NCOR2, CD24, NANOG, SOX2, POU5F1; NCOR2 with CD24, NANOG, SOX2, POU5F1 (**Fig. 1H**). TAM-treated patients with primary resistance showed a positive correlation with HDAC1 and NCOR2, while ESR1 correlated positively with NANOG, SOX2, POU5F1 and negatively with CD44. Finally, TAM-treated patients with secondary resistance HDAC1 displayed a positive correlation with ESR1, NCOR2, CD24, NANOG, SOX2 and POU5F1; ESR1 with NCOR2, CD24, NANOG, SOX2 and POU5F1; NCOR2 with NANOG, SOX2 and POU5F1.

### 3.2 TAM differentially modulates clonogenicity and stemness based on the exposure model

We found that 0.2 μM TAM and upward did show significant colonicity reduction (**Fig. 2A**). Conversely, TAM concentrations from 0.1 μM and above caused a significant decrease in mammosphere formation compared to the control group. 0.1 μM TAM was selected as the starting point because it was the lowest concentration at which we did not observe a reduction in clonogenicity without more than 20% reduction in mammosphere formation.

**Fig. 2.**
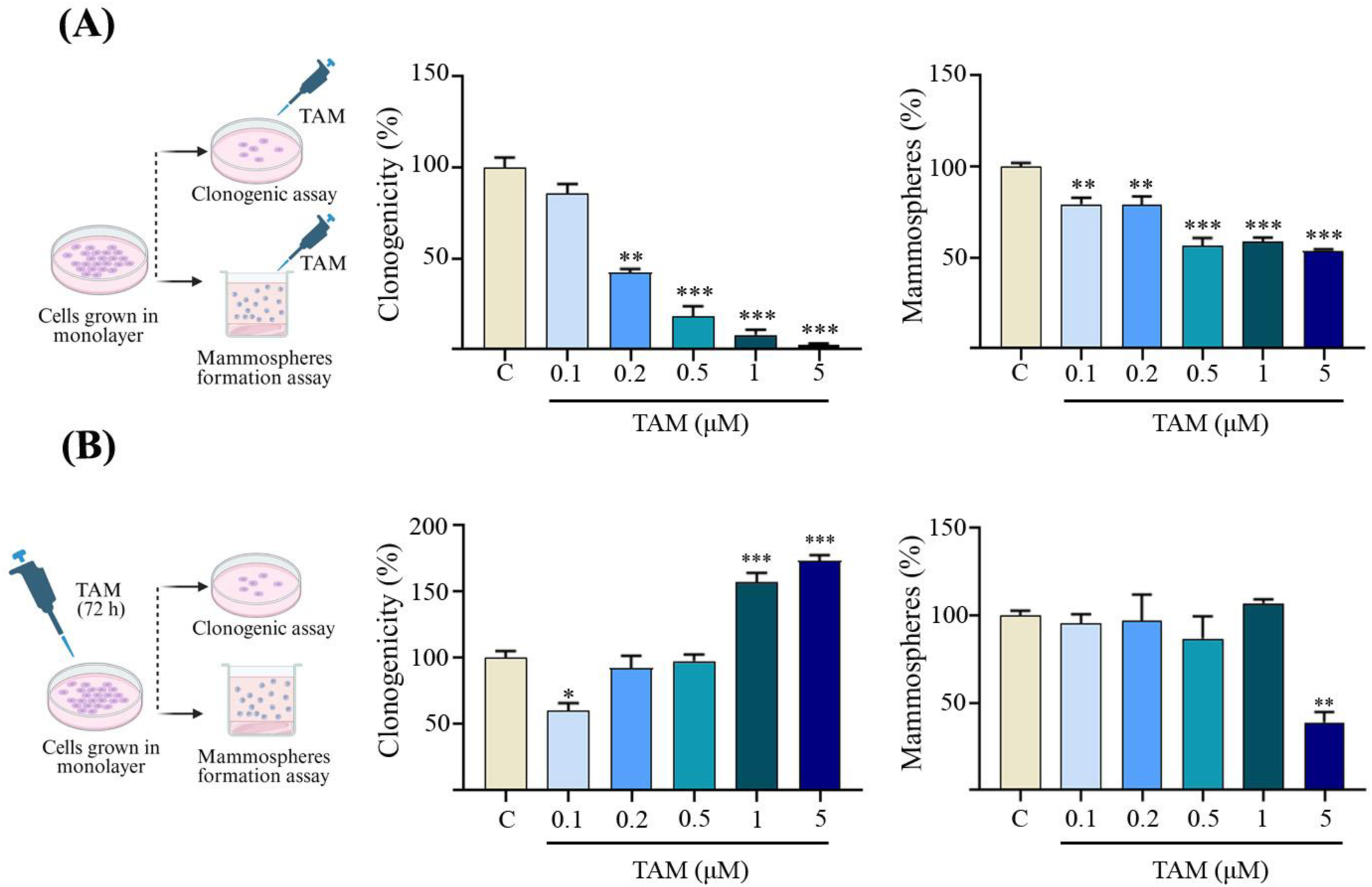
TAM differentially modulates clonogenicity and stemness based on the exposure model. **(A)-** MCF-7 cells were seeded on adherent plates for clonogenic assay or ultra-low adherent plates for mammospheres formation assay and treated with TAM (0.1- 5 μM) or vehicle (C). (\*\**p*<0.01, \*\*\**p*<0.001 vs. C). **(B)-** MCF-7 cells were grown in monolayer and treated with TAM (0.1- 5 μM) or vehicle (C) with μM for 72 h. Then, the trypsinized cells were grown on adherent plates for clonogenic assay or ultra-low adherent plates for mammosphere formation assay. (\*\**p*<0.01, \*\*\**p*<0.001 vs. C). All data in each graph are expressed as mean values ± SEM of three independent experiments. Experimental data sets were compared by One-way ANOVA and Dunnett’s Multiple Comparison post hoc test).

Moreover, given that our model is based on pretreatment of MCF-7 cells with CPF to the experiments, we decided to perform clonogenic and mammosphere formation assay cells that were pretreated for 72 h to different concentrations of TAM (**Fig. 2B**). TAM at 0.1 μM induced a reduction of the clonogenic capacity (40±5.8% below C, *p*<0.05), whereas cells exposed to 0.2 μM and 0.5 μM of TAM showed no difference to the control group, and cells treated with 1 and 5 μM of TAM experienced a significant increase in clonogenicity (57.3±7% and 73.7±7.1% over C, *p*<0.001, respectively). In the mammosphere formation assay, merely 5 μM TAM-pretreated exhibited a significant reduction of capacity to form mammospheres (61.03±5.86% below C, *p*<0.05). Given that exposure to 0.2 μM and 0.5 μM TAM did not significantly alter either clonogenic capacity nor the ability to form mammospheres, they were selected for the upcoming experiments.

### 3.3 CPF and TAM co-pretreatment affects clonogenicity and stem-like features in MCF-7 cells

To determine if CPF exposure affects TAM clonogenicity response and/or stemness selection, we used the *CPF pretreatment model.* We found that 50 μM CPF-pretreated MCF-7 cells showed an increase in the clonogenicity (73.2±14.76 over C, *p*<0.001). After seeding, the addition of 0.1 μM TAM caused a reduction of the clonogenic capacity of cells exposed to high concentrations of CPF (*p*<0.001 vs 50 μM CPF). We found an increase in the capability to form mammospheres induced by 0.05 μM CPF (32.3±3.31% over C, *p*<0.05). Nonetheless, when comparing C+0.1 μM TAM vs. 0.05 μM CPF+0.1 μM TAM we noticed that the cells exposed to the toxicant display a greater capacity to form mammospheres, even when it was administered concomitantly with the drug (*p*<0.01 vs C+TAM 0.1 μM) (**Fig. 3A**).

**Fig. 3.**
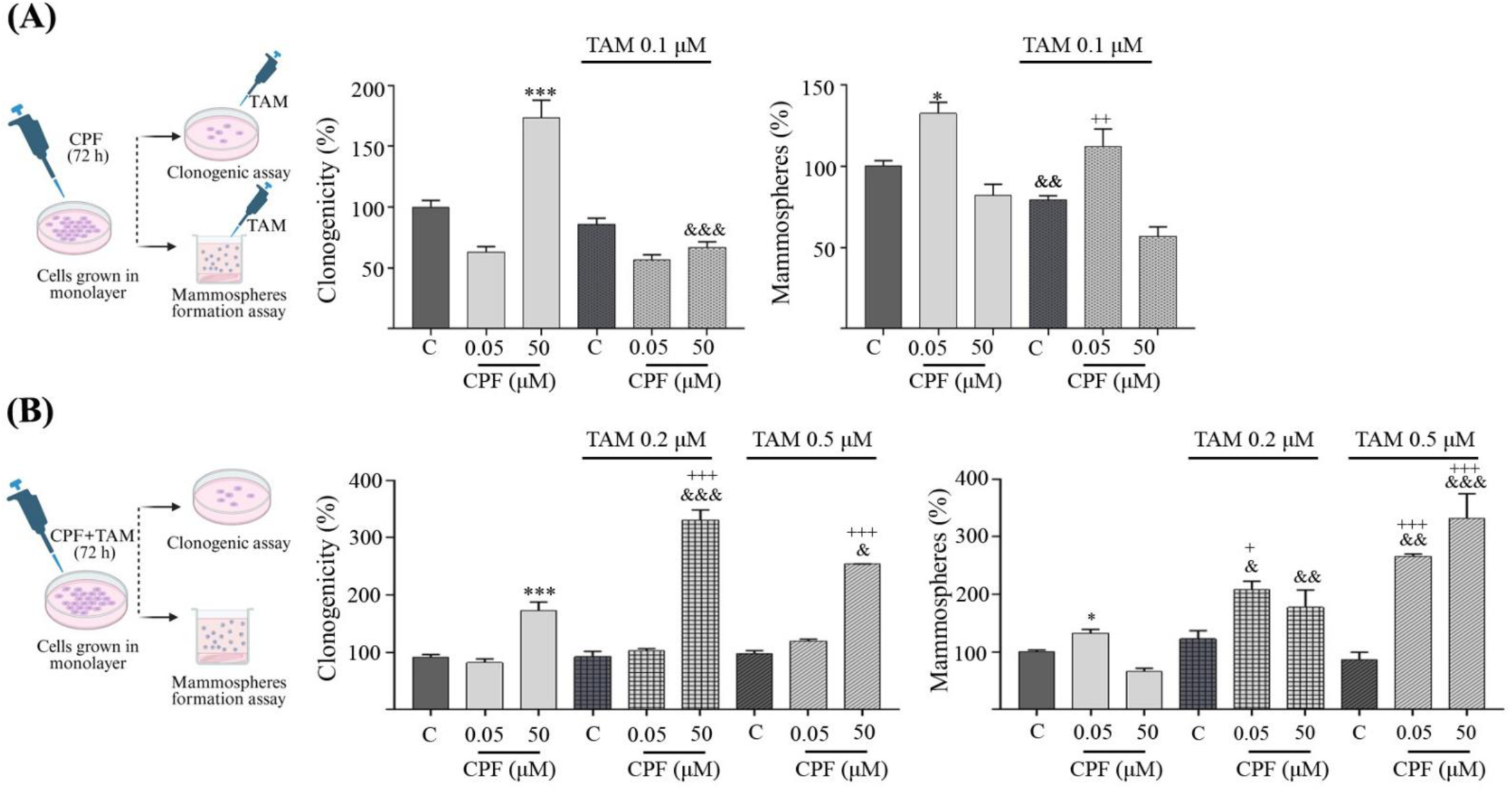
CPF and TAM co-treatment affects clonogenicity and stem-like features in MCF-7 cells. **A-** *CPF pretreated model.* MCF-7 cells were treated with CPF (0.05 and 50 μM) or vehicle (C) for 72 h. Then, the trypsinized cells were grown on adherent plates for clonogenic assay or ultra-low adherent plates for mammospheres formation assay. Finally, they were treated with 0.1 μM TAM. (\**p*<0.05, \*\*\**p*<0,001 vs. C; ^&&^*p*<0.01, ^&&&^*p*<0.001 vs. 0.05 μM or 50 μM CPF; ^++^*p*<0.01 vs C+0.1 μM TAM). **(B)-** *CPF+TAM concurrently co- pretreated model.* MCF-7 cells were treated with CPF (0.05 and 50 μM) or vehicle (C) with or without 0,2 μM or 0,5 μM TAM for 72 h. Then, the trypsinized cells were grown on adherent plates for clonogenic assay or ultra- low adherent plates for mammospheres formation assay. (\**p*<0.05, \*\*\**p*<0,001 vs. C; ^&^*p*<0.05, ^&&^*p*<0.01, ^&&&^*p*<0.001 vs 0.5 μM or 50 μM CPF; ^+^*p*<0.05, ^+++^*p*<0.01 vs C+0.2 μM TAM or C+ 0.5 μM TAM). All data in each graph are expressed as mean values ± SEM of three independent experiments. Experimental data sets were compared by Two-way ANOVA and Tukey’s Multiple Comparison post hoc test.

To assess whether simultaneous exposure to CPF and TAM is able to induce changes in clonogenicity and/or stemness phenotype, the *CPF+TAM concurrently co-pretreated model* was used. In this model, we noticed that the rise in clonogenic capacity in the pretreated MCF-7 cells to 50 μM CPF (*p*<0.001 vs. C) is exacerbated with the synchronous presence of 0.2 μM TAM (*p*<0.001 vs. CPF 50 μM) y 0.5 μM TAM (*p*<0.05 vs. CPF 50 μM). We have demonstrated that the increase in subpopulation of cells with stemness properties induced by 0.05 μM CPF (32.3±3.31% over C, *p*<0.05) is heightened when the toxicant is administered simultaneously with with 0.2 μM TAM (*p*<0.05 vs. CPF 0.05 μM) and 0.5 μM TAM (*p*<0.01 vs. CPF 0.05 μM). Remarkably we found that the concurrent administration of 50 μM CPF with both concentrations of TAM increases the quantity of formed mammospheres (*p*<0.01 vs. 50 μM CPF+0.2 μM TAM, *p*<0.001 vs. 50 μM CPF+0.5 μM TAM), albeit the sole exposure to the toxicant does not raise CSC (**Fig. 3B**).

### 3.4 In silico docking of CPF and TAM could interact with the ligand binding domain of the oestrogen receptor

The *in silico* models show that the 3 ligands: E2, CPF and TAM bind with favorable energy values to the same binding site in the receptor, corresponding to the one described as LBD. For CPF, the most favourable pose with binding energy values of -6.2 kcal/mol seems to be stabilized by hydrogen bonds with Thr347 and probably His524, and hydrophobic interactions with Leu391 and Phe404 (the latter by stacking interaction) (**Fig. 4A**). On the other hand, the most favorable pose for TAM (-9.5 kcal/mol) seems to occur by a salt bridge with Asp351 and multiple hydrophobic interactions, with Leu346, Ala350, Trp383, Leu387, Leu391, Phe404 and Leu428 (**Fig. 4B**). **Fig. 4C** shows that both CPF and TAM could interact with residues in the endogenous ligand-binding domain, oestradiol, of the oestrogen receptor.

**Fig. 4.**
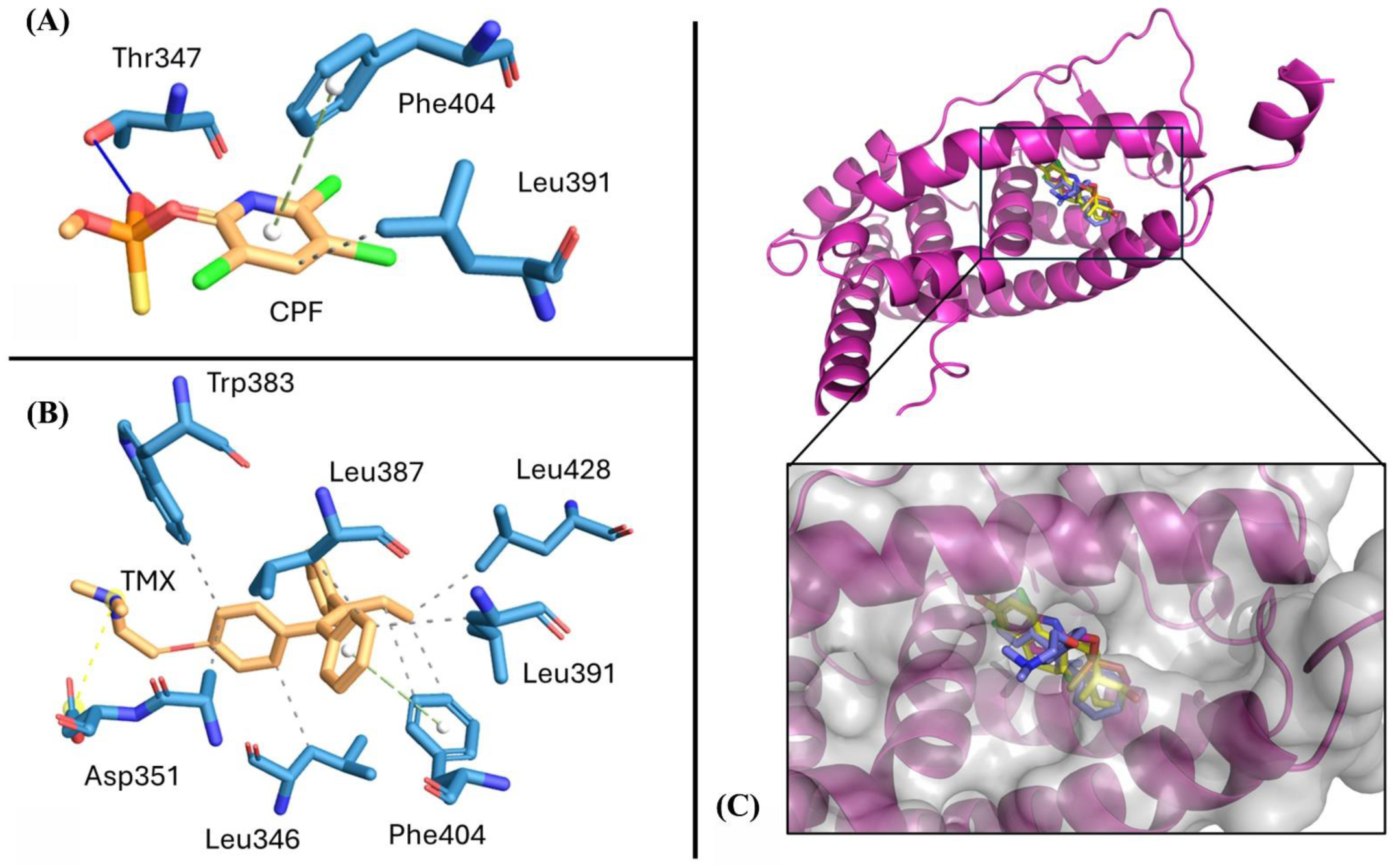
In silico docking of CPF and TAM could interact with the ligand binding domain of the oestrogen receptor. Main interactions between the LBD and **(A)-** CPF, and **(B)-** TMX. Solid blue lines: hydrogen bonds; green dashed lines: stacking hydrophobic interactions; green dotted lines: hydrophobic interactions; yellow dotted lines: salt bridges. **(C)-** Overall view of the whole hERα (magenta), and detailed view of the LBD showing the superposed ligands: estradiol (yellow), CPF (dark pink), TAM (blue). A transparent surface is shown to depict the narrowness of the binding site.

### 3.5 CPF and TAM co-pretreatment modulates stemness markers in monolayer-cultured MCF-7 cells

The assessment of stemness and resistance to anti-estrogen therapy markers was performed using the *CPF+TAM concurrently co-pretreated model*, as the most substantial differences in clonogenicity and mammosphere formation were observed under these conditions. We found that both CPF concentrations induce a decrease in the percentage of CD24^+^/CD44^+^ cells (12.69±0.67, 9.7±1.7 below C, *p*<0.01). Complementarily, we observed that exposure to TAM induced a significant decrease in this cell subpopulation (16.83±0.12 below C, p<0.001) and that co-pretreatment with 0.05 μM CPF + 0.5 μM TAM induced a significant increase when compared to exposure to TAM alone (*p*<0.01) (**Fig. 5A**).

**Fig. 5.**
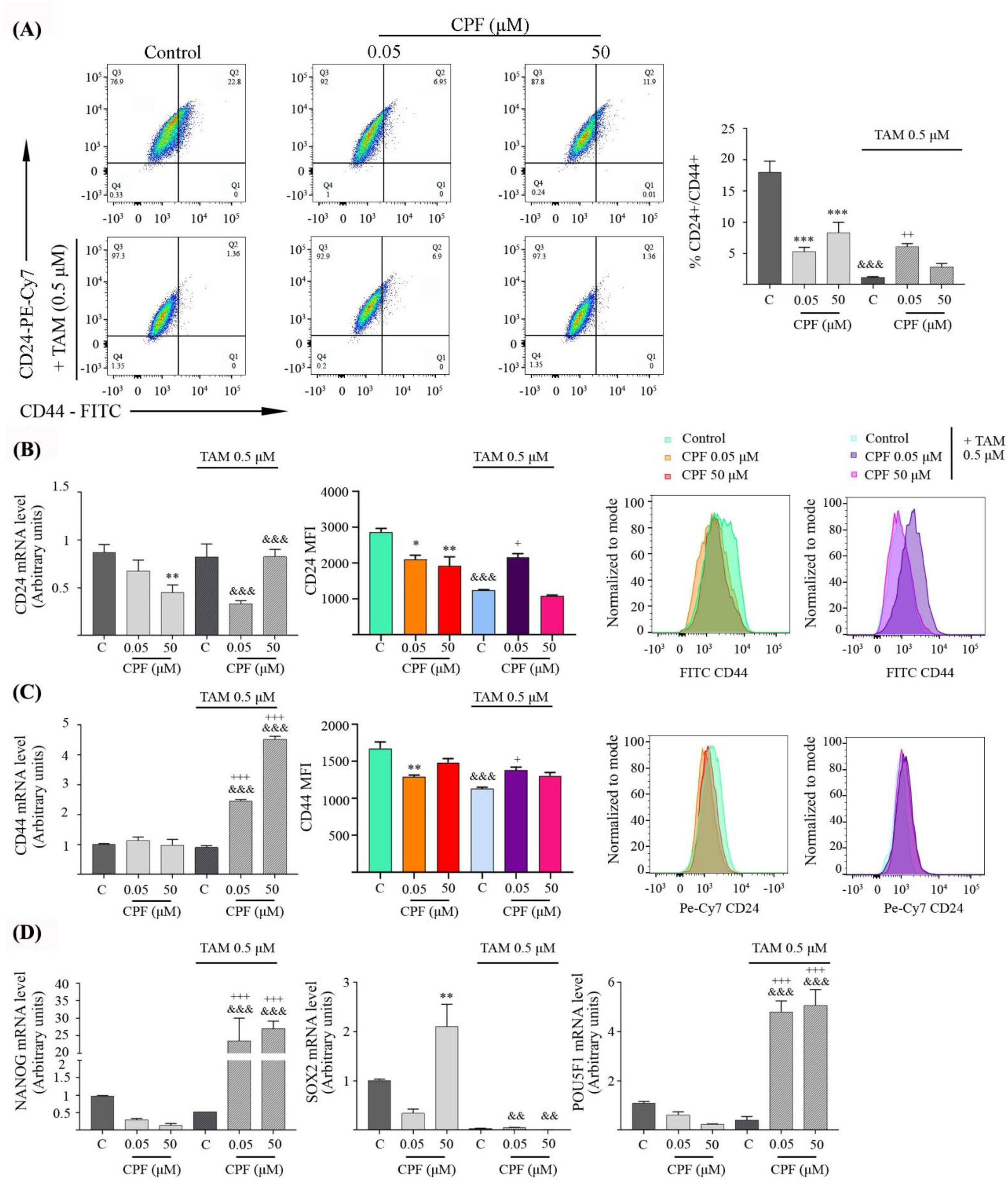
CPF and TAM co-pretreatment modulates stemness markers in monolayer-cultured MCF-7 cells. **(A)-** MCF-7 cells grown in monolayer were exposed to CPF (0.05 and 50 μM) or vehicle (C) with or without 0.5 μM TAM for 72 h. Flow cytometry analysis of the expression of CD44 and CD24 was performed. Cells were double stained with anti-CD44-FITC and anti-CD24-PE-Cy7. The percentage of CD24^+^/CD44^+^ positive cells was determined. (\*\*\**p*<0,001 vs. C; ^&&&^*p*<0.001 vs. C; ^++^*p*<0.01, vs C+TAM 0.5 μM). **(B)-** Relative mRNA levels of CD24 were determined by RT-qPCR. The results were normalized with respect to β-actin mRNA expression. (\*\**p*<0,01 vs. C; ^&&&^*p*<0.001 vs. 0.05 or 50 μM CPF). The level of CD24 expression was calculated by determination of the MFI. (\**p*<0,05, \*\**p*<0,01 vs. C; ^&&&^*p*<0.001 vs. C; ^+^*p*<0.05, vs. C+0.5 μM TAM). **(C)-** Relative mRNA levels of CD44 were determined by RT-qPCR. The results were normalized with respect to β-actin mRNA expression. (^&&&^*p*<0.001 vs. 0.05 or 50 CPF μM; ^+++^*p*<0.001 vs. C+0.5 μM TAM). The level of CD44 expression was calculated by determination of the MFI. (\*\**p*<0,01 vs. C; ^&&&^*p*<0.001 vs. C; ^++^*p*<0.01, vs. C+0.5 μM TAM). **(D)-** MCF-7 cells grown in monolayer were exposed to CPF (0.05 and 50 μM) or vehicle (C) with or without 0,5 μM for 72 h. Relative mRNA levels of NANOG, SOX2 and POU5F1 were determined by RT-qPCR. The results were normalized with respect to β-actin mRNA expression. (\**p*<0.05, \*\*\**p*<0,001 vs. C; ^&^*p*<0.05, ^&&&^*p*<0.001 vs. 0.5 μM or 50 μM CPF; ^++^*p*<0.01, ^+++^*p*<0.001 vs. C+0.5 μM TAM). All data in each graph are expressed as mean values ± SEM of three independent experiments. Experimental data sets were compared by Two-way ANOVA and Tukey’s Multiple Comparison post hoc test.

Complementally, 50 μM CPF induced a significant reduction in CD24 mRNA (0.42±0.08 above C, *p*<0.01). Cells co-pretreated with 0.05 μM CPF and 0.5 μM TAM showed decreased CD24 expression levels (*p*<0.001) and those co-pretreated with 50 μM CPF and 0.5 μM TAM showed increased mRNA expression levels for CD24 (*p*<0.05) when compared to CPF exposure alone. When analysing the MFI of CD24 we observed that cells co-pretreated with both pesticide concentrations show a decrease in expression levels (CPF 0.05 μM: 762±123.1 below C*, p*<0.05; 50 μM CPF: 935±252 below C, *p*<0.01), as well as those exposed to 0.5 μM TAM (1618±23.38 below C, *p*<0.001). We found that co- pretreatment with 0.05 μM CPF+0.5 μM TAM induced an increase in CD24 compared to C+0.5 μM TAM (*p*<0.05) (**Fig. 5B**).

In monolayer-cultivated cells we found that cells with concurrent exposure to both concentrations of CPF and 0.5 μM TAM experienced a rise in CD44 mRNA levels when compared to cells exposed to CPF alone (*p*<0.001). We observed that 0.05 μM CPF induced a decrease in CD44 expression levels when determining MFI (384±20.92 below C, *p*<0.001). Additionally, TAM induced a decrease in the expression of this cell surface marker (547±20.21 below C, *p*<0.001). In MCF-7 cells simultaneously exposed to 0.05 μM CPF and 0.5 μM TAM we observed a significant increase of CD44 when compared to cells exposed to TAM alone (*p*<0.05) (**Fig. 5C**).

Finally, as shown in **Fig. 5D**, we analysed the dedifferentiation markers NANOG, SOX2 and POU5F1, which codes for the OCT4 protein. Cells co-pretreated with 50 μM CPF showed increased levels of SOX2 mRNA expression (1.1±0.44 over C, *p*<0.01). Cells co-pretreated with 0.05 μM CPF and 0.5 μM TAM showed an increase in NANOG and POU5F1 (*p*<0.001), as did those treated with 50 μM CPF and 0.5 μM TAM (*p*<0.001) when compared with CPF alone. Those cells co-pretreated with vehicle or 50 μM CPF and 0.5 μM TAM showed a significant decrease in SOX2 levels (*p*<0.01 vs. C or 50 μM CPF).

### 3.6 CPF and TAM co-pretreatment orchestrates the regulation of ET resistance markers in monolayer-cultured MCF-7 cells

The exposure of MCF-7 cells to low concentrations of CPF induced a decline in mRNA expression of ESR1 and HDAC (0.57±0.01, 0.4±0.03 below C, respectively, *p*<0.05) while rising levels of NCOR2 (0.97±0.4 over C, *p*<0.05) (**Fig. 6A**). Conversely, the treatment with 50 μM CPF prompted a reduction of NCOR2 levels (0.7±0.04 below C, *p*<0.05). Cells which underwent concurrent co-pretreatment with 0.05 μM CPF and 0.5 μM TAM exhibited higher expression levels of mRNA of ESR1 (*p*<0.001), HDAC1 (*p*<0.05) and NCOR2 (*p*<0.001) and simultaneous administration of 50 μM CPF and 0.5 μM TAM promoted an increment in ESR1 (*p*<0.001), HDAC1 (*p*<0.001) and NCOR2 (*p*<0.001) when compared with CPF alone.

**Fig 6.**
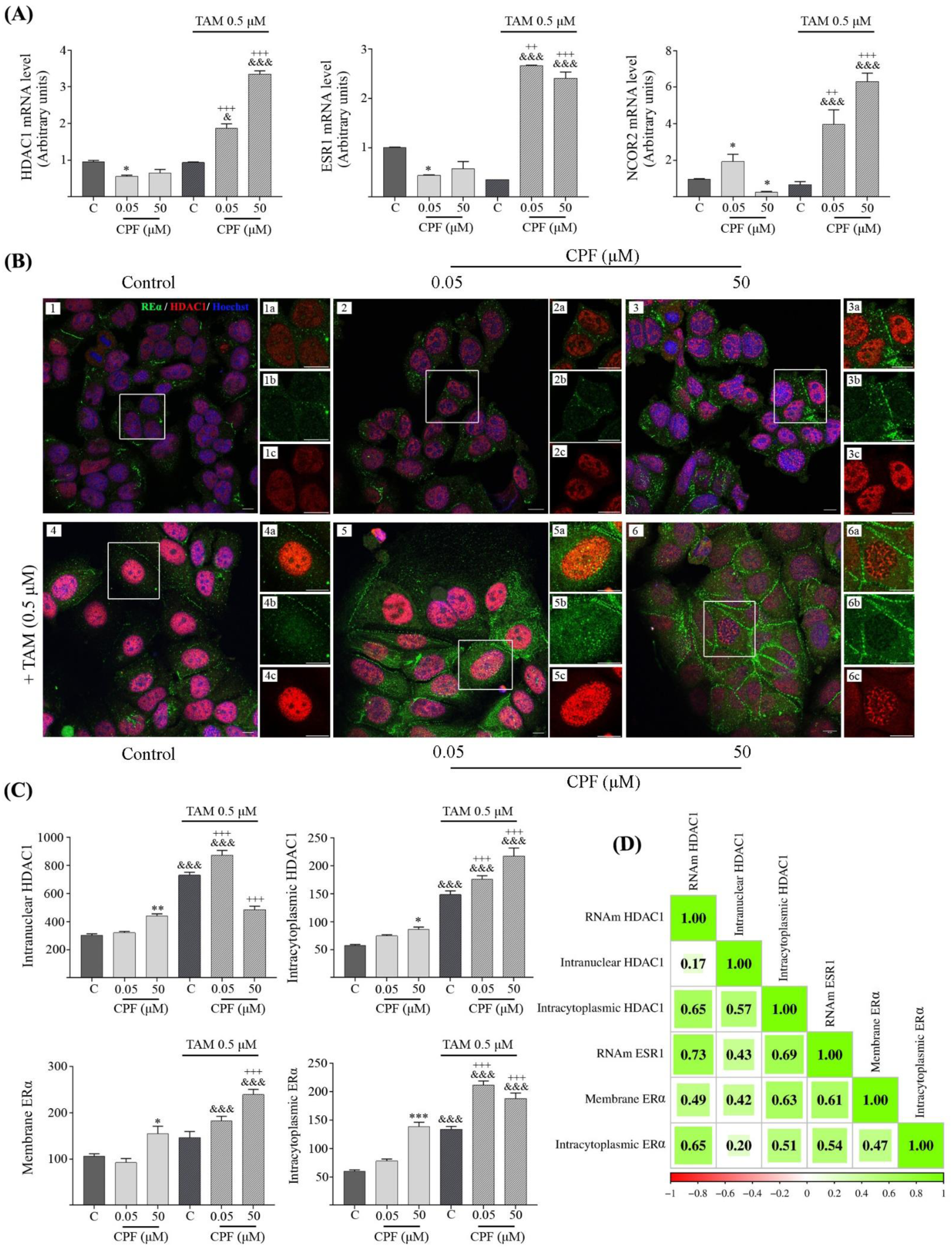
CPF and TAM co-pretreatment orchestrates the regulation of ET resistance markers in monolayer- cultured MCF-7 cells. **A-** MCF-7 cells grown in monolayer were exposed to CPF (0.05 and 50 μM) or vehicle (C) with or without 0.5 μM TAM for 72 h. Relative mRNA levels of HDAC1, ESR1 and SMRT mRNA were determined by RT-qPCR. The results were normalized with respect to β-actin mRNA expression. (\**p*<0.05, \*\*\**p*<0,001 vs. C; ^&^*p*<0.05, ^&&&^*p*<0.001 vs 0.5 μM or 50 μM CPF; ^++^*p*<0.01, ^+++^*p*<0.001 vs C+ 0.5 μM TAM). **(B)-**Images of MCF-7 grown in monolayer treated con CPF (0.05 and 50 μM) or vehicule (C) with or without 0.5 μM TAM for 72 hs with Hoechst staining **(1,6)**, immunofluorescence for ERα **(1b, 6b)**, HDAC1 **(1c, 6c)**, and merge **(1, 6 and 1a, 6a)** were taken on a Zeiss LSM880 confocal microscope. Bar scale: 10 μm. Magnification: 63x **(1,6)** and digital Zoom **(1a-b-c, 6a-b-c). (C)-** The level of HDAC1 (intranuclear and intracytoplasmic) and ER (intracytoplasmic and membrane) expression was calculated by determination of the MFI in the Z-project of each of the image stacks. (\**p*<0.05, \*\*\**p*<0,001 vs. C; ^&&&^*p*<0.001 vs CPF 0.5 μM or 50 μM; ^+++^*p*<0.001 vs C+TAM 0.5 μM. **D) -** Correlation matrix plot with lower triangle color intensity and size of the square proportional to the correlation coefficients. The color scale bar ranged from −1.0 (red) to 1.0 (green). The size and color of the square indicate the magnitude of the correlation between parameters. Red and green, respectively, denote negative and positive correlations. All data in each graph are expressed as mean values ± SEM of three independent experiments. Experimental data sets were compared by Two-way ANOVA and Tukey’s Multiple Comparison post hoc test.

The cells that were co-pretreated with 50 μM CPF showed increased intranuclear HDAC1 expression (135±9.34 over C, *p*<0.01). The same effect was observed when cells were co-pretreated with TAM in both, those exposed to vehicle (426.5±9.35 over C, *p*<0.05) and those exposed to 0.05 μM CPF (*p*<0.001, 0.05 μM CPF vs. 0.05 μM CPF+0.5 μM TAM). Additionally we found an increase in intranuclear HDAC1 expression levels in those cells co-pretreated with 0.05 μM CPF and 0.5 μM TAM (*p*<0.001) and a decrease in HDAC1 levels in cells co-pretreated with 50 μM CPF and 0.5 μM TAM *(p*<0.001) relative to those treated with vehicle and TAM (**Fig. 6B-C**).

A striking finding was the increase in intracytoplasmic HDAC1 expression observed with cells co- pretreated with 50 μM CPF (29.15±2.65 over C, *p*<0.05). TAM co-pretreatment induced an increase in intracytoplasmic HDAC1 expression in both those exposed to vehicle (*p*<0.001 vs. C) and those exposed to 0.05 μM CPF (*p*<0.001 vs. 0.05 μM CPF) and 50 μM CPF (*p*<0.001 vs. 50 μM CPF). We found an increase in intracytoplasmic HDAC1 expression levels in those cells co-pretreated with 0.05 μM CPF and 0.5 μM TAM (*p*<0.001) as well as those pretreated with 50 μM CPF and 0.5 μM TAM (*p*<0.001) with respect to those treated with vehicle and TAM 0.5 μM.

Furthermore, 50 μM CPF increased membrane expression of ERα (48.9±5.3 over C*, p*<0.05). We observed that co-pretreatment with TAM raised membrane cell ERα expression in cells exposed to 0.05 μM CPF (*p*<0.001 vs. 0.05 μM CPF) and 50 μM CPF (*p*<0.001 vs. 50 μM CPF). Additionally, we found an increase in membrane ERα levels in those cells co-pretreated with 50 μM CPF and 0.5 μM TAM, with respect to those treated with vehicle and 0.5 μM TAM (*p*<0.001).

Cells pretreated with 50 μM CPF also showed increased levels of ERα expression in the cytoplasm (78.5±2.29 over C, *p*<0.001). Additionally, co-pretreatment with TAM induced an increase in intracytoplasmic ERα expression in both those exposed to vehicle (*p*<0.001 vs. C) and those exposed to 0.05 μM CPF (*p*<0.001 vs. 0.05 μM CPF) and 50 μM CPF (*p*<0.001 vs 50 μM CPF). Finally, we found an increase in the expression levels of ERα intracytoplasmic in those cells co-pretreated with 0.05 μM CPF and 0.05 μM TAM (*p*<0.001) as well as in those co-pretreated with 50 μMCPF and 0.5 μM TAM (*p*<0.001) with respect to those treated with vehicle and 0.5 μM TAM.

Additionally, we assessed the presence of correlation between ESR1 and HDAC mRNA levels, as well as the levels of both proteins in the different cellular compartments (**Fig 6D**). We found that HDAC1 mRNA correlates positively with intracytoplasmic HDAC1 levels, ESR1 mRNA as well as with membrane and intracytoplasmic ERα levels. There is a positive correlation between nuclear and intracytoplasmic HDAC1 levels. Intranuclear HDAC1 correlates positively with ESR1 mRNA and ERα membrane expression. Intracytoplasmic HDAC1 correlated positively with both ESR1 mRNA and membrane and intracytoplasmic ERα expression. ESR1 mRNA correlates positively with membrane (*r^2^*=0.61, *p*=0.001) and intracytoplasmic ERα levels. There is a positive correlation between membrane and intracytoplasmic expressed ERα levels.

### 3.7 CPF and TAM co-treatment induces context-dependent correlation patterns between endocrine resistance and stemness markers in MCF-7 cells

In mammospheres formed by MCF-7 cells pretreated with CPF 0.05 μM we observed decreased levels of CD24 (0.46±0.1 below C, *p*<0.01), along with higher levels of CD44 (5.75±1.25 over C, *p*<0.01) and POU5F1 (1.06±015 over C, *p*<0.001) (**Fig. 7A**). Those cells which were exposed to 50 μM CPF showed increased expression of POU5F1 (1.54±0.43 over C, *p*<0.01) and NANOG (3.96±0.88 over C, *p*<0.05). Additionally, mammospheres obtained from cells with co-pretreatment with 0.05 μM CPF and 0.5 μM TAM displayed raised levels of CD24 mRNA (*p*<0.001 vs. 0.05 μM CPF) and lower levels of CD44 (*p*<0.001). Those co-pretreated with 50 μM CPF and 0.5 μM TAM expressed greater levels of CD24 (*p*<0.001), POU5F1 (*p*<0.05) and NANOG (*p*<0.05) when compared to cells solely exposed to CPF 50 μM.

**Fig. 7.**
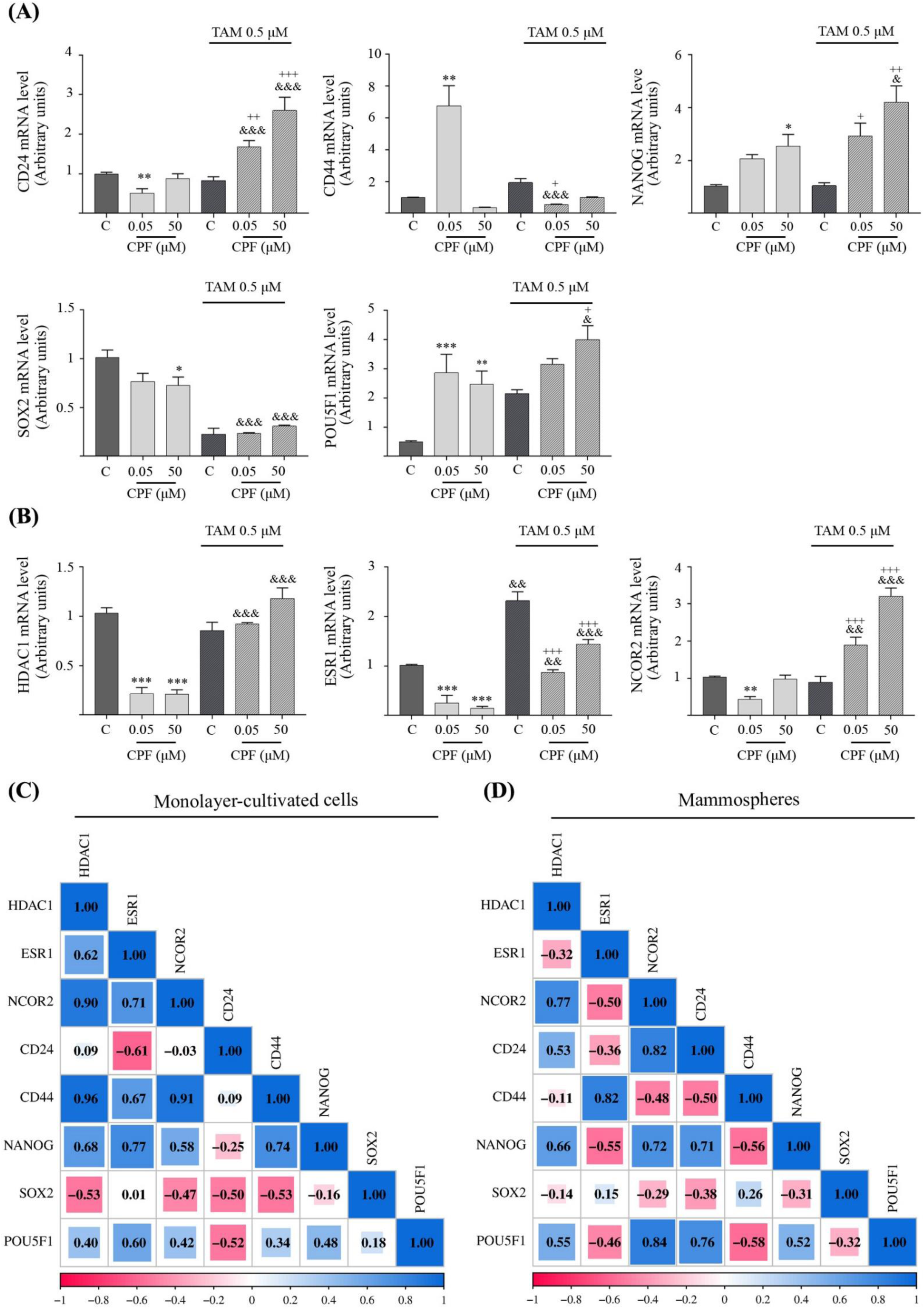
CPF and TAM co-treatment induces context-dependent correlation patterns between endocrine resistance and stemness markers in MCF-7 cells. Relative mRNA levels of CSC markers: CD24, CD44, NANOG, SOX2 and POU5F1 **(A)** and ET resistance markers: HDAC1, ESR1 and SMRT **(B)** were determined by RT-qPCR in the mammospheres obtained from MCF-7 cells co-pretreated for 72 hours with CPF (0.05 and 50 μM) or vehicle (C) with or without 0.5 μM TAM. The results were normalized with respect to β-actin mRNA expression. (\**p*<0.05, \*\**p*<0.01, \*\*\**p*<0,001 vs. C; ^&^*p*<0.05, ^&&^*p*<0.01, ^&&&^*p*<0.001 vs CPF 0.5 μM or 50 μM; ^+^*p*<0.05, ^++^*p*<0.01, ^+++^*p*<0.001 vs C+TAM 0.5 μM). **(C-D)-** Correlation matrix plot with lower triangle color intensity and size of the square proportional to the correlation coefficients. The color scale bar ranged from −1.0 (pink) to 1.0 (blue). The size and color of the square indicate the magnitude of the correlation between parameters. Pink and blue, respectively, denote negative and positive correlations. All data in each graph are expressed as mean values ± SEM of three independent experiments. Experimental data sets were compared by Two-way ANOVA and Tukey’s Multiple Comparison post hoc test.

When analyzing markers of ET resistance (**Fig. 7B**), we noticed that mammospheres obtained from co-pretreated cells to either of the concentrations of CPF displayed lower levels of mRNA of ESR1 (CPF 0.05 μM = 0.76±0.15 below C, CPF 50 μM = 0.87±0.04 below C, *p*<0.001) and HDAC1 (CPF 0.05 μM = 0.84±0.11 below C, CPF 50 μM = 0.84±0.04 below C, *p*<0.001). Notably only those co-pretreated with CPF 0.05 μM showed decreased levels of NCOR2 (0.6±0.07 below C, *p*<0.01). Mammospheres from MCF-7 cells co-pretreated with TAM and either concentration of CPF exhibited a significant increment in levels of mRNA of ESR1 (*p*<0.01) and HDAC (*p*<0.001) when compared to those from cells with sole treatment with the toxicant. Furthermore, the levels of ESR1 and HDAC1 with both concentrations of CPF were proximate to the levels found in the control mammospheres. Lastly, we noticed that mammospheres formed from cells co-pretreated with TAM and both concentrations of CPF showed significant increase in NCOR2 mRNA (*p*<0.001 vs. 0.05 CPF or 50 μM CPF).

In MCF-7 cells grown in monolayer that received co-pretreated with CPF+TAM, several markers were found to have strong positives such as HDAC1 with ESR1, NCOR2, CD44, NANOG and POU5F1. Additionally, ESR1 correlates positively with NCOR2, CD44, NANOG and POU5F1 as well as NCOR2 with CD44, NANOG and POU5F1. Finally, CD44 presents strong positive correlations with NANOG and POU5F1 and NANOG with POU5F1. Conversely, CD24 shows negative correlations with ERS1 and SOX2 consistently shows negative correlations with HDAC1, NCOR2, CD44 and NANOG (**Fig. 7C**).

In mammospheres obtained from MCF-7 cells co-pretreated with CPF+TAM we observed the strong positive correlation between HDAC with NCOR2, CD24, NANOG and POU5F1 as well al NCOR2 with CD24, NANOG and POU5F1, CD24 with NANOG and NANOG with POU5F1. In contrast, we observed a strong negative correlation among ERS1 with NCOR2, NANOG and POU5F1, NCOR2 with CD44, CD24 with CD44 and CD44 with NANOG and POU5F1 (**Fig. 7D**).

## 4. Discussion

Breast cancer has remained one of the most commonly diagnosed types of malignant tumors in recent years, placing first in worldwide frequency in 2020 and second in 2022, and continues to be the number one cause of cancer death in women as well as the fourth one overall (World Health Organization, 2024). Although patients with HR-positive breast cancers generally respond to ET pproximately 30% develop resistance, which compromises long-term survival (Aggelis & Johnston, 2019; Nolan et al., 2023). Among the proposed risk factors, it is increasingly suggested that environmental pollutants are able to modulate signaling pathways involved in the emergence and progression of metastatic tumor cells (Koual et al., 2020). Chlorpyrifos (CPF) is a highly lipophilic pesticide, which has raised concerns about its persistence in the environment and its bioaccumulation. In Europe, its agricultural use has been banned since 2020, and in 2025 it was added to the Stockholm Convention as a persistent organic pollutant (European Commission, 2025). In the US, in 2021, the EPA banned its use on crops intended for human and animal consumption, but this regulation was overturned by a federal court, restoring the previous tolerances for CPF use, which led the EPA to issue a new regulation maintaining 11 specific uses on fruits and vegetables such as apples, blueberries, soybeans and sugar beets, among others (United States Environmental Protection Agency, 2025). In Argentina, its commercialization and fractionation have been banned since 2023 (SENASA, 2021).

Given that the well-documented role as an endocrine disruptor (Abd-Elhakim et al., 2021; Fouyet et al., 2022; Ventura et al., 2016), its capacity to induce epithelial-to-mesenchymal transition (EMT) (Lasagna et al., 2020, 2022), and its ability to expand subpopulations of cells with stem cell-like features (Lasagna et al., 2025), we deemed it essential to investigate its potential contribution to the development of resistance to ET in breast cancer. Therefore, we set out to investigate whether CPF can modulate the response to TAM in an ER+ *in vitro* breast cancer model, as well as its influence on molecular markers associated with endocrine resistance and stemness phenotype, two features strongly linked to poor prognosis and disease progression, as evidenced by our analysis of public breast cancer patient datasets.

Initially, we observed that many GO analysis include three categories: BP (TGF- β, regulation of epithelial cell proliferation and response to steroid hormone), CC (cell-cell junction, actin-based cell projection) and MF (Actin and β-catenin binding) as well as the differentially expressed signaling pathways (MAPK, PI3K-AKT and Wnt) in TAM-resistant MCF-7 cells are modulated by CPF, all of which prompted us to further study the participation of CPF in the development of TAM resistance. Additionally, we noticed that patients who were not selected for TAM treatment showed higher levels of HDAC1 and lower levels of ESR1, as well as a more undifferentiated profile with lower levels of CD24 and higher levels of CD44, SOX2 and NANOG.

Historically, HDACs have been investigated as potential prognostic factors for breast cancer. Therefore, we analyzed public databases and found that low levels of HDAC1 expression are positively correlated with shorter disease-free survival in breast cancer patients who received TAM. In addition, patients who showed primary resistance to endocrine therapy had lower levels of HDAC1 compared to those who did not relapse or who experienced secondary endocrine resistance. This discovery is consistent with that reported by Seo et al (2014), who found that high expression of HDAC1 was positively correlated with good overall survival in the HR-positive group (Seo et al., 2014). These findings were surprising, as it has been postulated that HDAC1 and HDAC2 increase chemosensitivity to anticancer drugs by inhibiting the expression of multidrug resistance-related genes in colon cancer models (Xu et al., 2012). Thus, HDAC inhibitors increased TAM efficacy by downregulating the expression of the receptor tyrosine kinase RET (rearranged during transfection), which is a promising candidate related to recovery TAM sensitivity (Griseri et al., 2016).

In patients treated with TAM we observed a positive correlation between CD24 and other stemness- associated markers such as NANOG, SOX2, and POU5F1. In contrast, we found that subjects who were not selected for TAM treatment exhibited a strong correlation among HDAC1, ESR1, and NCOR2, as well as with stemness-related markers. In addition, the information collected from patients included in public databases, who did not relapse and of those with primary endocrine resistance, showed that HDAC1 expression levels were only positively correlated with NCOR2, but not with dedifferentiation markers. Furthermore, we observed that data from patients who developed secondary endocrine resistance showed a very similar correlation pattern to patients who had not been selected for TAM treatment; moreover, in the latter subgroup we found a positive correlation of HDAC1 with ESR1 and NCOR2, as well as with the dedifferentiation markers CD24, NANOG, SOX2 and POU5F1. The aforementioned results suggest that the selected markers, as well as the correlations among them, could potentially serve as predictors of response to anti-oestrogen therapy.

Regarding the TAM response, in the pretreatment model, we noticed a reduction of the anti-estrogenic effects. TAM at 1 and 5 uM concentrations increases clonogenicity, demonstrating that the drug exerts a selection effect on cells and that those that survive have greater proliferative capacity. Nevertheless, 5 μM TAM diminishes mammosphere formation as expected. Simoes et al (2015) showed that short- term treatment with 1 μM of the anti-estrogens 4OH-TAM decreases cell proliferation but increases BCSC activity through JAG1-NOTCH4 receptor activation both in patient-derived samples and xenograft (PDX) tumors. In our experiments, we used growth factors deprived-serum. The smaller and fewer mammospheres in absence of E2 could explain this effect as previously reported (Y. Zhang et al., 2012).

In the CPF pretreated model we observed that CPF does not modify the response of cells to TAM. However, in the CPF+TAM concurrently co-pretreated model, cells which survived the exposure show greater clonogenicity and enrichment of stem cell-like cells. In our previous work, we demonstrated that the use of a SERDs called ICI 182,780 -also known as fulvestrant-, which binds to the ERα with high affinity and leads to its proteasomal degradation, counteracts the decreased the number and size of mammospheres formed (Lasagna et al., 2022). In this regard, it is also known that inhibition of ERα with its inverse agonist, XCT-790, or its knockdown in breast cancer cells significantly reduces the size and efficiency of mammosphere formation (Muduli et al., 2023). Therefore, the presence of the ERα receptor is mandatory for CPF to increase its ability to select cells with a more aggressive phenotype in the presence of TAM. This finding could be related to the results of the *in silico* analysis suggesting that a simultaneous binding of more than one ligand, whether TAM, CPF or E2, to the ERα seems unlikely. However, depending on the concentrations *in vivo*, and the relative affinities of each ligand, it could be possible the occurrence of a competitive displacement of E2 by both CPF and TAM, considering that both molecules contain residues that can interact with the receptor.

For a better characterization of the cells that survive co-pretreatment with CPF+TAM, we evaluated their expression of endocrine resistance and stemness markers. We recently reported that low concentrations of CPF induced an increase in the percentage of CD24^+^/CD44^+^ cells in a triple- negative model (Lasagna et al., 2025). Here, we found that the pesticide decreased this subpopulation in monolayer-cultured MCF-7 cells with a reduction in CD24 mRNA levels and protein expression. Basal-like breast cancer subtype is usually enriched in CD24^-/low^ cells, which, in turn, are less frequent in luminal tumors (Taurin & Alkhalifa, 2020). Therefore our findings are suggestive that exposure to CPF generates a phenotypic shift towards more aggressive and antiestrogenic therapy resistance tumors.

Strikingly, co-pretreatment with 0.05 uM CPF+TAM induces an increase in CD24^+^/CD44^+^ subpopulation, a reduction in mRNA levels and an increment in CD24 protein levels when compared to exposure to TAM alone. Regarding CD44 expression we observed that co-treatment with both concentrations of CPF+TAM increases CD44 mRNA expression levels, finding which does not fully correlate with protein expression levels. Although initially Al-Hajji et al. (2003) demonstrated that CD24^-/low^/CD44^high^ cells display tumorigenic properties, recent findings show that the subpopulation of CD24^-/low^/CD44^-/low^ cells, which are enriched in the luminal cell lines such as T47D, MCF-7 and BT- 474, possess metastatic and tumorigenic properties (Vikram et al., 2020). In the mammospheres obtained from 0.05 uM CPF-pretreated cells, we found a reduction in the expression levels of CD24 and an increase in CD44, configuring a stem-like phenotype similar to that described by Al-Hajji et al (2003). Conversely, when analyzing the effect of co-treatment with CPF+TAM, we observed an increase in CD24 and a decrease in CD44 expression levels. In accordance with this, we have previously reported that CPF induced an increment of CD24 mRNA levels in a triple negative tumor model (Lasagna et al., 2025).

In both the mammospheres obtained from cells pretreated with CPF and in the MCF-7 cells cultured in monolayers as well as the mammospheres that received pretreatment with CPF+TAM, we observed higher levels of NANOG and POU5F1 mRNA and reduced levels of SOX2. The importance of NANOG in the maintenance of pluripotency and proliferation is well documented and up-regulation of its expression is critically related to tumourigenic cells in cancer (C. Zhang et al., 2016). Vikram et al (2020) evaluated NANOG expression in CD24^-/low^/CD44^-/low^ cells of luminal cell lines and CD24^-/low^/CD44^high^ cells of the highly invasive triple-negative cell line MDA-MB-231 and found similar expression levels, indicating its possible key role in the maintenance of pluripotency in these cell populations. It is known that embryonic stem cell genes such as MYC, NANOG, and OCT4 are known to be overexpressed in the poorly differentiated basal subtype of breast cancer (Taurin & Alkhalifa, 2020) and therefore our findings could be linked to the generation of more basal and, therefore, more aggressive phenotypes, lacking response to anti-estrogen therapy.

Co-pretreatment with CPF+TAM of monolayer-cultured cells increases HDAC1 protein expression (intranuclear and cytoplasmic), as well as HDAC1 and NCOR mRNA levels. These effects could prevent the transcriptional machinery from accessing the ESR1 gene promoters, thereby suppressing its transcriptional activity (Seto & Yoshida, 2014). Therefore, the increase in both clonogenicity and enrichment in the stemness phenotype subpopulation associated with the activation of transcriptional repression mechanisms observed in cells exposed to pretreatment with CPF+TAM, could be explained by an acquisition of a more aggressive basal-like phenotype, whose proliferative capacity has become independent of ERα action. Furthermore, HDAC1 is known to be capable of silencing multiple epithelial genes, giving rise to cells with hybrid epithelial-mesenchymal phenotypes. Intermediate states of the mesenchymal transformation became the focus of the investigations because of the capacity of the plasticity of these subpopulations. Thus, it has been demonstrated that this deacetylase is recruited to the CDH1 promoter and represses Slug-mediated E-cadherin expression. In this context, we have previously reported that CPF increases SLUG levels, reduces E- cadherin expression, and promotes a mesenchymal phenotype (Adhikary et al., 2014; Lei et al., 2010). Therefore, HDAC1 may represent a mediator of CPF-induced TEM, suggesting a new mechanism involved in the advanced stages of carcinogenesis triggered by environmental factors.

A surprising finding was the increase in HDAC1 expression at the intracellular level, since the expression of class I HDACs has been commonly found in the nucleus. It is known that this enzyme can also produce direct deacetylation of cytoplasmic proteins, regulating their activity (Dunaway & Pollock, 2022). In this regard, although most studies show that HDAC1 has antiangiogenic actions, a recent study has demonstrated that HDAC1 participates in proangiogenic mechanisms in the cytosol independently of transcription modulation. Although the molecular targets involved are not fully understood, we highlight this study because it suggests that the subcellular localization of HDAC1 generates a differential modulation of the processes in which this enzyme participates (Bazou et al., 2016).

Additionally, co-pretreatment with CPF+TAM increased the expression of ESR1 mRNA levels and ERα protein not only in the intracellular compartment but also in the external cell membrane. Initially, we were struck by the upward regulation of both ER and the HDAC1-NCOR2 repressor complex. However, Ghosh et al (Ghosh et al., 2018) have reported that the persistent organochlorine pesticide endosulfan in the MCF-7 cell line induces ERα overexpression associated with a significant increase in total HDAC activity and an increase in the expression levels of HDAC 1 and 3, suggesting that this mechanism could be an effect shared with other agents that act as endocrine disruptors. Complementary, and although it is widely accepted that ERα is predominantly observed in the nucleus, the existence of extranuclear estrogenic actions was reported more than 50 years ago. It is known that ERα signaling initiated in the membrane is capable of rapidly activating various signaling pathways such as tyrosine-protein kinase c-SRC, MAPK/ERK, and PI3K/AKT (Adlanmerini et al., 2022). We have demonstrated in previous studies that all these pathways are activated by CPF and are closely related to different stages of breast carcinogenesis, in this particular case to the development of resistance to antiestrogen therapy (Lasagna et al., 2022).

In monolayer cultures, we observed a positive correlation between endocrine therapy resistance markers, as well as between NANOG and POU5F1, both markers associated with the stem phenotype. This correlation pattern is consistent with the transcriptomic profiles identified in public databases of luminal breast cancer patients with secondary resistance to ET. Therefore, it is likely that CPF promotes the development of resistance to ET, and these correlation patterns could become biomarkers capable of predicting response to therapy, demonstrating the translational relevance of our work. It should be noted that, in our in vitro model, unlike the trends observed in clinical datasets, CD44 also showed a positive correlation with resistance markers, suggesting that it could represent a biomarker unique to the action of the toxin and its ability to induce pluripotency. In contrast, mammospheres showed a markedly different correlation profile: NCOR2 correlated negatively with ESR1 and CD44, and we observed a negative correlation between CD44 and CD24. Furthermore, the dedifferentiation markers NANOG and POU5F1 correlated negatively with ESR1 and CD44. These findings suggest that CPF and TAM co-pretreatment modulates resistance and stemness markers in a microenvironment- and architecture-dependent manner, emphasizing the importance of using 2D and 3D culture models when investigating endocrine resistance mechanisms.

## 5. Conclusions

In summary, we can conclude that the simultaneous exposure to the endocrine disruptor CPF and the antiestrogen agent TAM led to a selection of a subpopulation of cells that exhibit increased clonogenicity, enhanced mammosphere-forming ability, and upregulates the stemness markers NANOG and POU5F1. This subpopulation also shows higher levels of mRNA and elevated protein expression of REα and HDAC1. The role of the significance of the CD24/CD44 rate remains controversial and has shown a dependence on the microenvironment. Thus, co-pretreatment induces an increase in CD44 and a decrease in CD24 in cells grown in monolayer, while in mammospheres, the opposite effect was observed. Finally, we identify a resistance and stemness marker signature induced by CPF+TAM that closely resembles the profile observed in the dataset of patients who acquired tamoxifen resistance. All of these findings indicate that co-pretreatment with CPF+TAM induces a more aggressive, undifferentiated and basal-like cell phenotype, and therefore, a loss of response to the antiestrogenic action of TAM. These results highlight the translational relevance of our work, as they provide evidence regarding the different mechanisms by which CPF could induce resistance to endocrine therapy in luminal breast cancer. Our findings support existing bans on CPF in several countries by providing biological plausibility for its potential role in cancer progression.

---

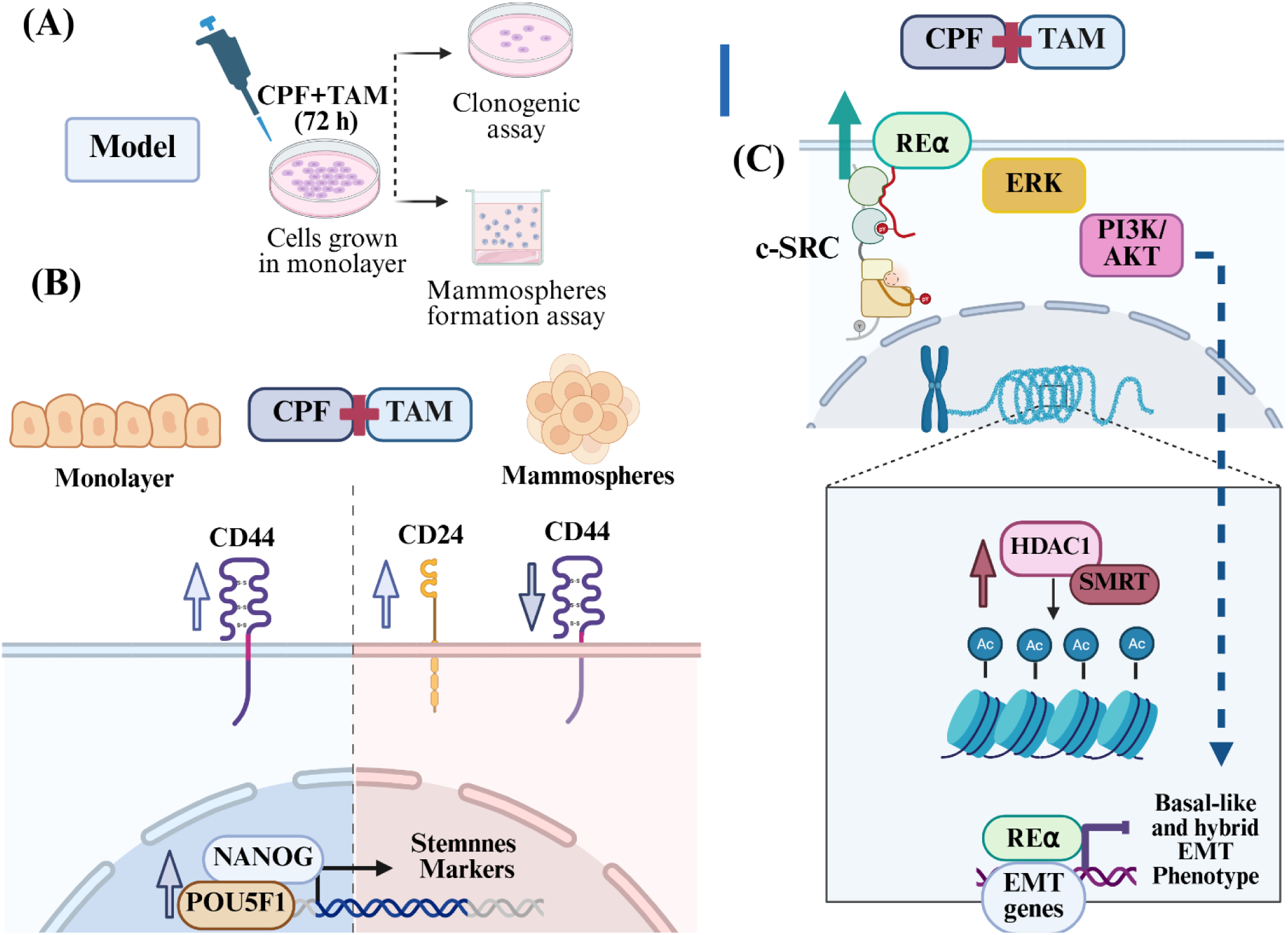
Graphical abstract. **(A)-** CPF+TAM Concurrently Co-Pretreated Model. **(B)-** Effect of the CPF+TAM Concurrently Co-Pretreated Model on Stemness Markers. **(C)-** Effect of the CPF+TAM Concurrently Co- Pretreated Model on Anti-Estrogen Therapy Resistance Markers. Created in https://BioRender.com.

